# Hierarchical effects of choice-related activity and neural encoding during feature selective attention

**DOI:** 10.1101/2020.03.10.986349

**Authors:** Jennifer L. Mohn, Joshua D. Downer, Kevin N. O’Connor, Jeffrey S. Johnson, Mitchell L. Sutter

## Abstract

Selective attention is necessary to sift through, form a coherent percept of, and make behavioral decisions on the vast amount of information present in most sensory environments. How and where selective attention is employed in cortex and how this perceptual information then informs the relevant behavioral decisions is still not well understood. Studies probing selective attention and decision making in visual cortex have been enlightening as to how sensory attention might work in that modality; whether or not similar mechanisms are employed in auditory attention is not yet clear. Therefore, we trained rhesus macaques on a feature selective attention task, where they switched between reporting changes in temporal (amplitude modulation, AM) and spectral (carrier bandwidth) features of a broadband noise stimulus. We investigated how the encoding of these features by single neurons in primary (A1) and secondary (lateral belt, ML) auditory cortex were affected by the different attention conditions. We found that neurons in A1 and ML showed mixed-selectivity to the sound and task features. We found no difference in AM encoding between the attention conditions. We found that choice-related activity in both A1 and ML neurons shifts between attentional conditions. This finding suggests that choice-related activity in auditory cortex does not simply reflect motor preparation or action, and supports the relationship between reported choice-related activity and the decision and perceptual process.

**New & Noteworthy:** We recorded from primary and secondary auditory cortex while monkeys performed a non-spatial feature attention task. Both areas exhibited rate-based choice-related activity. The manifestation of choice-related activity was attention-dependent, suggesting that choice-related activity in auditory cortex does not simply reflect arousal or motor influences, but relates to the specific perceptual choice. The lack of temporal-based choice activity is consistent with growing evidence that subcortical, but not cortical, single neurons inform decisions through temporal envelope following.

## Introduction

The auditory system is often faced with the difficult challenge of encoding a specific sound in a noisy environment, such as following a conversation in a loud room. The neural mechanisms by which the auditory system attends to one sound source and ignores distracting sounds are not yet understood. Studies probing the mechanisms underlying auditory attention in cortex have been largely concerned with task engagement, wherein the effects of active performance on neural activity is compared to those of passive listening. Studies in auditory cortex (AC) utilizing this paradigm have shown that task engagement can improve behaviorally-relevant neural sound discrimination (Atiani et al. 2014; Bagur et al. 2018; Buran et al. 2014; Carcea et al. 2017; Francis et al. 2018a; Niwa et al. 2012a, 2015; von Trapp et al. 2016), modulate neuronal tuning (Fritz et al. 2003, 2007; Fritz 2005; Lee and Middlebrooks 2011; Lin et al. 2019; Yin et al. 2014), alter the structure of correlated variability within neural populations (Downer et al. 2015, 2017a), and more (Massoudi et al. 2014; Angeloni and Geffen 2018; Osmanski and Wang 2015; Sutter and Shamma 2011). Though informative, this active/passive paradigm makes it difficult to disentangle arousal and motor effects from the mechanisms more specifically employed in selectively attending to a single sound source or feature amidst auditory ‘clutter’.

Studies of the neural basis of auditory selective attention at the single neuron level are rare (Schwartz and David 2018), and non-spatial auditory feature-selective attention has been relatively unexplored (Downer et al. 2017b). Feature-selective attention, which segregates particular sound features, such as intensity or fundamental frequency, is essential for tasks such as discriminating between talkers in a noisy environment (Bregman 1994; McDermott 2009; Bizley and Cohen 2013; Shinn-Cunningham 2008; Woods and McDermott 2015). Furthermore, it can prove useful for listeners to switch between attended sound features because the most distinctive feature dimensions may vary across sources (Woods and McDermott 2015; Bregman 1994).

In visual cortex, feature-based attention has been suggested to follow a gain model similar to spatial attention, where responsivity to the attended feature increases in cells tuned to the attended feature and decreases in cells tuned to orthogonal features (Martinez-Trujillo and Treue 2004; Maunsell and Treue 2006). Studies of spatial attention in AC single neurons suggest that AC employs a mechanism similar to that reported in visual cortex, where a gain in neural activity increases when attention is directed into the receptive field of a neuron and, conversely, gain decreases when attention is directed outside the receptive field (Engle and Recanzone 2013; Lee and Middlebrooks 2011; Scott et al. 2007). We endeavored to see if feature-selective attention in AC is also facilitated by a gain in activity in neurons tuned to an attended feature.

How and where task relevant sensory information is transformed into a decision in the brain is still largely unclear. There have been mixed reports of activity correlated to the reported decision in AC (Christison-Lagay et al. 2017; Elgueda et al. 2019; Guo et al. 2019; Niwa et al. 2012b; Runyan et al. 2017; Tsunada et al. 2016; Tsunada and Cohen 2014). This choice-related activity has been reported in some studies as early as primary auditory cortex (A1) (Atiani et al. 2014; Bathellier et al. 2012; Bizley et al. 2013; Christison-Lagay et al. 2017; Christison-Lagay and Cohen 2018; Francis et al. 2018b, 2018b; Gronskaya and von der Behrens 2019; Huang et al. 2019; Niwa et al. 2012b). As one progresses further along the auditory cortical hierarchy, there is either an increasingly larger proportion of neurons showing activity correlated to the decision, or the nature of the choice signal changes (Atiani et al. 2014; Niwa et al. 2013; Tsunada et al. 2016). Both cases suggest that the sensory evidence informing task-relevant decisions is transformed as the information moves up the processing stream (Bizley and Cohen 2013; Hackett 2011; Huang and Brosch 2020; Romanski et al. 1999).

There has also been uncertainty as to whether the reported choice activity in AC could be more reflective of motor influences than perceptual or decision-related influences. Go/No-Go tasks are typically used in auditory cortical studies, and these tasks require movement for report of one choice, but not the other (Brosch 2005; Niwa et al. 2013); forced-choice tasks reduce this uncertainty by requiring movements for either report (Guo et al. 2019). It has been well documented that movement can modulate auditory cortical activity (Eliades and Wang 2003; Guo et al. 2019; Schneider et al. 2014). Here, we employ a Yes/No forced-choice task format in which a movement is required for both responses in order to disentangle motor-related from choice-related activity in AC.

We investigated whether a mechanism for feature-selective attention similar to feature-based attention in visual cortex is employed in primary (A1) and secondary (middle lateral belt, ML) auditory cortex using noise that was amplitude modulated (AM) or bandwidth restricted (ΔBW). Monkeys were presented sounds that varied either in spectral (ΔBW) or temporal (AM) dimensions, or both, and performed a detection task in which they reported change along one of these feature dimensions. In this study, we focus on the amplitude modulation feature, as it has been well studied and is a salient communicative sound feature for humans and other animals (Schnupp 2006; Shannon et al. 1995; Van Tasell et al. 1987; Wang et al. 2007) and can be helpful in sound source segregation (Bregman 1994; Grimault et al. 2002). Spectral content changes were used as a difficulty-matched attentional control. We hypothesized we would see a gain in AM encoding when animals were cued to attend to that feature, compared to when they were cued to attend ΔBW changes. We also examined choice-related activity in AC, hypothesizing to find a larger proportion of neurons with significant choice-related activity in higher-order AC (ML) than in A1.

## Materials and Methods

### Subjects

Subjects were two adult rhesus macaques, one male (13kg, 14-16 years old), one female (7kg, 17-19 years old). All procedures were approved by the University of California-Davis Animal Care and Use Committee and met the requirements of the United States Public Health Service policy on experimental animal care.

### Stimuli

Stimuli were constructed from broadband Gaussian (white) noise bursts (400 ms; 5 ms cosine ramped), 9 octaves in width (40 to 20480 Hz). Four different seeds were used to create the carrier noise, which was frozen across trials. To introduce variance along spectral and temporal dimensions, the spectral bandwidth of the noise was narrowed (ΔBW) and/or the noise envelope was sinusoidally amplitude modulated (AM). The extent of variation in each dimension was manipulated to measure behavioral and neural responses above and below threshold for detecting each feature.

Sound generation methods have been previously reported (O’Connor et al., 2011). Briefly, sound signals were produced using an in-house MATLAB program and a digital-to-analog converter (Cambridge Electronic Design [CED] model 1401). Signals were attenuated (TDT Systems PA5, Leader LAT-45), amplified (RadioShack MPA-200), and presented from a single speaker (RadioShack PA-110) positioned approximately 1.5 m in front of the subject centered at the interaural midpoint. Sounds were generated at a 100 kHz sampling rate. Intensity was calibrated across all sounds (Bruel & Kjaer model 2231) to 65 dB at the outer ear. It is important to note that some methods of generating ΔBW introduce variation in that sound’s envelope, however we implemented a synthesis method that constructs noise using a single-frequency additive technique and thus avoids introducing envelope variations that could serve as cues for ΔBW (Strickland and Viemeister 1997).

### Recording procedures

Each animal was implanted with a head post centrally behind the brow ridge and a recording cylinder over an 18 mm craniotomy over the parietal lobe using aseptic surgical techniques (O’Connor et al. 2005). Placement of the craniotomy was based on stereotactic coordinates of auditory cortex to allow vertical access through parietal cortex to the superior temporal plane (Saleem and Logothetis 2007).

All recordings took place in a sound attenuating, foam-lined booth (IAC: 2.9×3.2×2 meters) while subjects sat in an acoustically transparent chair (Crist Instruments). Three quartz-coated tungsten microelectrodes (Thomas Recording, 1-2 MΩ; 0.35 mm horizontal spacing; variable, independently manipulated vertical spacing) were advanced vertically to the superior surface of the temporal lobe. Extracellular signals were amplified (AM Systems model 1800), bandpass filtered between 0.3 Hz and 10 kHz (Krohn-Hite 3382), and then converted to a digital signal at a 50 kHz sampling rate (CED model 1401). During electrode advancement, auditory responsive neurons were isolated by presenting various sounds while the subject sat passively. When at least one auditory responsive single unit was well isolated, we measured neural responses to the two features while the subjects sat passively awake. At least 10 repetitions of each of the following stimuli were presented: the unmodulated noise, each level of bandwidth restriction, and each of the possible AM test modulation frequencies (described below). We also measured pure tone and bandpass noise tuning to aid in distinguishing area boundaries.

After completing these tuning measures, experimental behavioral testing and recording began. When possible, tuning responses to the tested stimuli were again measured after task performance, to ensure stability of electrodes throughout the recording. Contributions of single units (SUs) to the signal were determined offline using principal components analysis-based spike sorting tools from Spike2 (CED). Spiking activity was at least 4-5 times the background noise level. Fewer than 0.1% of spike events assigned to single unit clusters fell within a 1 ms refractory period window. Only recordings in which neurons were well isolated for at least 180 trials within each condition were included in analysis here.

### Cortical field assessment

Recording locations were determined using both stereotactic coordinates (Martin and Bowden 1996) and established physiological measures (Merzenich and Brugge 1973; Rauschecker and Tian 2000; Tian and Rauschecker 2004). In each animal, we mapped characteristic frequency (CF) and sharpness of bandpass noise tuning to establish a topographic distribution of each. Tonotopic gradient reversal, BW distribution, spike latency and response robustness to pure tones was used to estimate the boundary between A1 and ML and assign single units to an area (Downer et al. 2017a; Niwa et al. 2015). Recordings were assigned to their putative cortical fields *post hoc* using recording location, tuning preferences, and latencies.

### Feature attention task

This feature attention task has been previously described in detail (Downer et al. 2017b). The subjects performed a change detection task in which only changes in the attended feature were relevant for the task. Subjects moved a joystick laterally to initiate a trial, wherein an initial sound (the S1, always the 9-octave-wide broadband, unmodulated noise) was presented, followed by a second sound (S2) after a 400ms inter-stimulus interval (ISI). The S2 could be identical to the S1, it could change by being amplitude-modulated (AM), it could change by being bandwidth restricted (ΔBW), or it could change along both feature dimensions.

Only three values of each feature (AM, ΔBW) were presented, limiting the size of the stimulus set in order to obtain reasonable power for neural data analysis. The stimulus space was further reduced by presenting only a subset of the possible co-varying stimuli. Within each recording session, we presented 13 total stimuli. To equilibrate difficulty between the two features, we presented values of each feature so that one was near threshold, one was slightly above, and one far above threshold.

Thresholds were determined for each feature independently for each subject using a range of six levels for each feature before three feature values for each animal were selected and the co-varying feature attention task began. For Monkey U, the ΔBW values were 0.375, 0.5, and 1 octave and the AM depth values were 28%, 40%, and 100%. For Monkey W, the ΔBW values were 0.5, 0.75, and 1.5 octaves and the AM depth values were 40%, 60%, and 100%.

For all analyses in which data are collapsed across subjects, ΔBW values and AM values are presented as ranks (ΔBW 0-3 and AM 0-3) (e.g., Fig. 1). Within a given session, AM sounds were presented at only a single modulation frequency. Across sessions, a small set of frequencies was used (15, 22, 30, 48, and 60 Hz). The AM frequency was selected randomly each day. Subjects were cued visually via an LED above the speaker as to which feature to attend (green or red light, counterbalanced between subjects). Additionally, each block began with 60 “instruction” trials in which the S2s presented were only altered along the attended feature dimension (i.e., sounds containing the distractor feature were not presented). Subjects were to respond with a “yes” (up or down joystick movement, counterbalanced across subjects) on any trial in which the attended feature was presented, otherwise, the correct response was “no” (opposite joystick movement). We chose upward or downward joystick movement to avoid influences on single neuron choice activity dependent on contralateral movements. Such movement related activity has been recently reported in other studies (Guo et al. 2019). Hits and correct rejections were rewarded with a drop of water or juice and misses and false alarms resulted in a penalty (3–5 s timeout).

**Figure 1:**
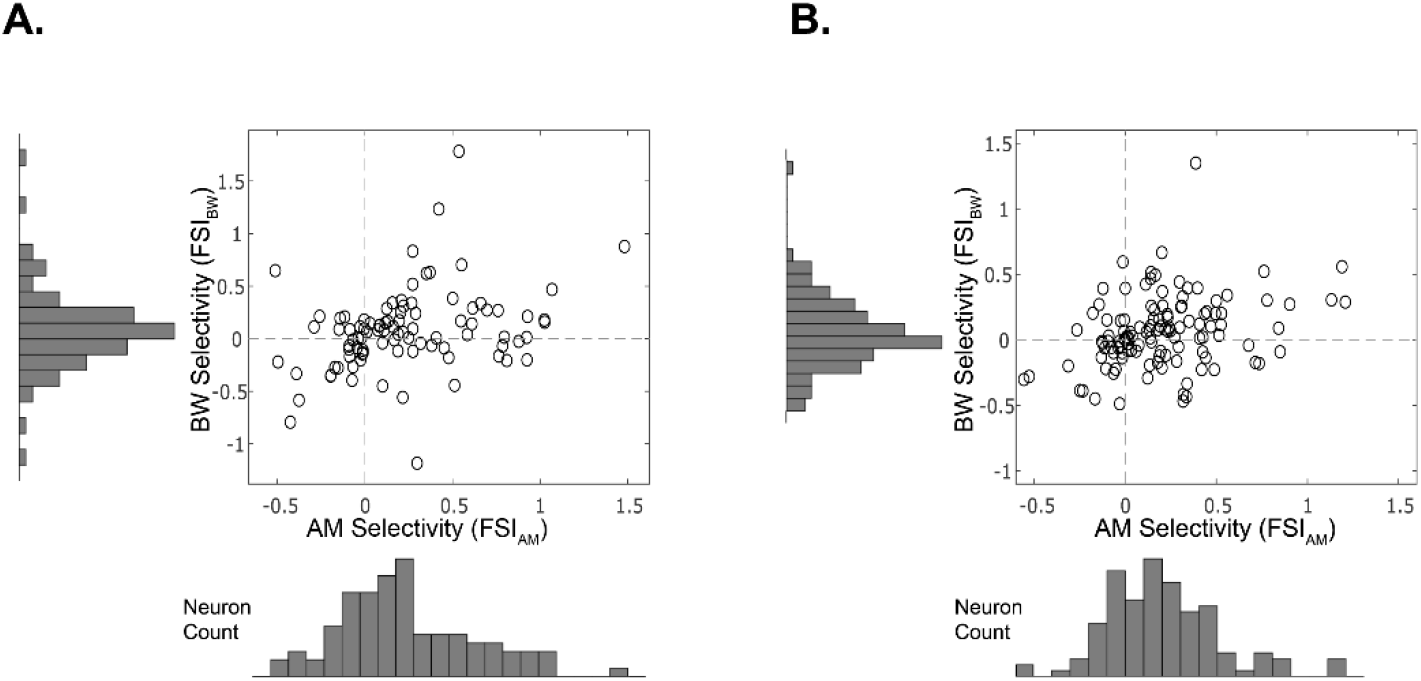
Single neuron feature selectivity index (FSI), a measure of how sensitive a neuron is to changes in each feature value separately. **A**. A1: a positive correlation between AM and ΔBW selectivity (Pierson rho = 0.3143, *p* = 0.002) **B.** ML: positive correlation between AM and BW selectivity (Pierson’s rho = 0.3109, *p* = 5.32 e-4)

During the test conditions, the S2 was unmodulated broadband noise (no change from S1) on 25% of the trials, co-varying on 25% of the trials, and contained only ΔBW or AM on 25% of the trials respectively. Sounds in the set were presented pseudo-randomly such that, over sets of 96 trials, the entire stimulus set was presented exhaustively (including all four random noise seeds). Block length was variable, based in part on subjects’ performance, to ensure a sufficient number of correct trials for each stimulus. Not including instruction trials, block length was at least 180 trials and at most 360 trials, to ensure that subjects performed in each attention condition at least once during the experiment. Subjects could perform each attention condition multiple times within a session. Only sessions in which subjects completed at least 180 trials per condition (excluding instruction trials) were considered for analysis in this study.

### Analysis of single neuron feature selectivity

Neurons’ firing rate responses across the range of values were calculated to derive a firing rate function for each feature. Functions were categorized based on whether firing rates increased as the level of the feature increased or decreased (‘increasing’ vs. ‘decreasing’ functions). Spike counts (SC) were calculated over the entire 400ms stimulus window. SCs in response to feature-present stimuli were normalized over the entire spike count distribution across both features, including unmodulated noise, for that cell. To characterize this response function, we calculated a feature-selectivity index (FSI) for each feature as follows:

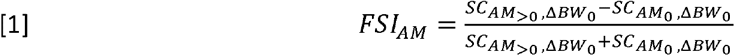

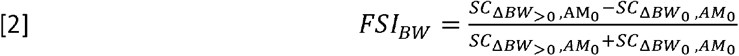

Where SC_x_ is the mean SC in response to the given set of stimuli designated by the subscript. A Kruskal-Wallis rank-sum test was preformed between distributions of SCs with the feature-present (feature level greater than 0) and those with the feature-absent (feature value of 0) to determine the significance of the FSI for each neuron. Cells that had a significant FSI for a given feature were categorized as encoding that feature.

### Phase projected vector strength

Vector strength (VS) is a metric that describes the degree to which the neural response is phase-locked to the stimulus (Goldberg and Brown 1969; Mardia and Jupp 2000). VS is defined as:

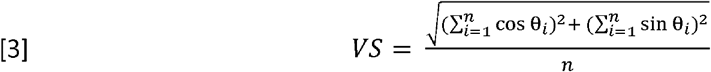

Where *n* is the number of spikes over all trials and θ_i_ is the phase of each spike, in radians, calculated by:

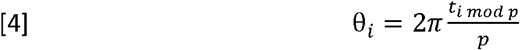

Where *t_i_* is the time of the spike (in ms) relative to the onset of the stimulus and *p* is the modulation period of the stimulus (in ms). When spike count is low, VS has a tendency to report as spuriously high. Phase projected Vector Strength (VS_pp_), is a variation on VS developed to help mitigate issues with low SC trials (Yin et al. 2011). VS_pp_ is calculated by first calculating VS for each trial, then the mean phase angle of each trial is compared to the mean phase angle of all trials, and the trial VS value is penalized if out of phase with the global mean response. VS_pp_ is defined as:

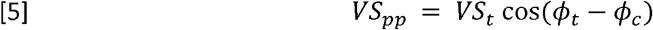

Where *VS_pp_* is the phase-projected vector strength per trial, VS_t_ is the vector strength per trial, as calculated in [1], and ϕ_*t*_ and ϕ_*c*_ are the trial-by-trial and mean phase angle in radians, respectively, calculated for each stimulus by:

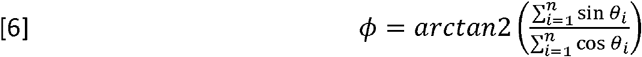

Where *n* is the number of spikes per trial (for ϕ_*t*_) or across all trials (for ϕ_*c*_). In this report, we use VS_pp_ exclusively to measure phase-locking, as SC tended to be relatively low and VS and VS_pp_ tend to be in good agreement with the exception of low SCs where VS_pp_ tends to be more accurate than VS (Yin et al. 2011). To determine significance of VS_pp_ encoding for each neuron, a Kruskal-Wallis rank-sum test was preformed between distributions of VS_pp_ values on trials with non-zero AM depths, to those from unmodulated noise trials. Of note, when we refer to the VS_pp_ in response to an unmodulated stimulus, this is a control measurement assuming the same modulation frequency as the corresponding AM frequency from that recording session.

### Analysis of neural discriminability

We applied the signal detection theory-based metric area under the receiver operating characteristic (ROCa) (Green and Swets 1974) to measure how well neurons could detect each feature. ROCa represents the probability an ideal observer can detect the presence of the target feature given only a measure of the neural responses (either firing rate or VS_pp_). To calculate ROCa, we partitioned the trial-by-trial neural responses into two distributions: those when the target feature was present in the stimulus and trials where it was absent. Then we determined the proportion of trials in each group where the neural response exceeds a criterion value. We repeated the measure using 100 criterion values, covering the whole range of responses. The plot of the probability of exceeding the criterion for feature-present trials (neural ‘hits’) versus the probability of exceeding the criteria for feature-absent trials (neural false alarms) plotted for all 100 criteria as separate points creates the ROC plot. The area under this curve is the ROCa. ROCa is bounded by 0 and 1, where both extremes indicate perfect discrimination between target feature-present and -absent stimuli, and 0.5 indicates a chance level of discrimination between the two distributions.

### Analysis of choice-related activity

Choice probability (CP) is an application of ROC analysis used to measure the difference between neural responses contingent on what the animal reports, for example, whether a stimulus feature is present or absent (Britten et al. 1992,1996). Similar to ROCa described above, CP values are bounded by 0 and 1, and a CP value of 0.5 indicates no difference (or perfect overlap) in the neural responses between ‘feature-present’ and ‘feature-absent’ reports. A CP value of 1 means for every trial that the animal reports a feature, the neuron fired more than on trials where the animal did not report the feature. A CP value of 0 means that, for every trial that the animal reports a feature, the neuron fired *less* than on trials where the animal did not report the feature. Stimuli that did not have at least 5 ‘yes’ and 5 ‘no’ responses were excluded from analyses. CP was calculated based on both firing rate and on VSpp. For rate-based CP, we calculated CP both for each stimulus separately, and pooled across stimuli. We calculated this stimulus-pooled CP by first separating the ‘yes’ and ‘no’ response trials within stimulus, then converting these rates into z-scores within a stimulus, then combined these z-scored responses across stimuli. This type of z-scoring has been found to be conservative in estimating CP (Kang and Maunsell 2012). CP was calculated during both the 400ms stimulus presentation (S2) and during the response window (RW), the time after stimulus offset and prior to the response (typically ~0.2 – 3s). The significance of each neuron’s CP was determined using a permutation test (Britten et al. 1996). The neural responses were pooled between the ‘feature-present report’ and ‘feature-absent report’ distributions and random samples were taken (without replacement). CP was then calculated from this randomly sampled set. This procedure was repeated 2000 times. The *p* value is the proportion of CP values from these randomly sampled repeats that were greater than the CP value from the non-shuffled distributions.

## Results

We recorded activity from 92 single units in A1 (57 from Monkey W, 35 from Monkey U) from 33 recording sessions and 122 single units in ML (49 from Monkey W, 73 from Monkey U) over 39 recording sessions as animals performed a feature-selective attention task.

### Feature tuning

There was no significant difference in the proportions of neurons in A1 and ML that encoded AM (47.8% A1, 38.5% ML; *p* = 0.08, χ2 test). We found a large proportion of neurons in both A1 and ML that were sensitive to the relatively small changes in ΔBW from the 9-octave wide unmodulated noise, though there was no difference in the proportion of ΔBW encoding neurons between areas (32.6% Al, 29.5% ML; *p* = 0.18, χ2 test) (Table 1).

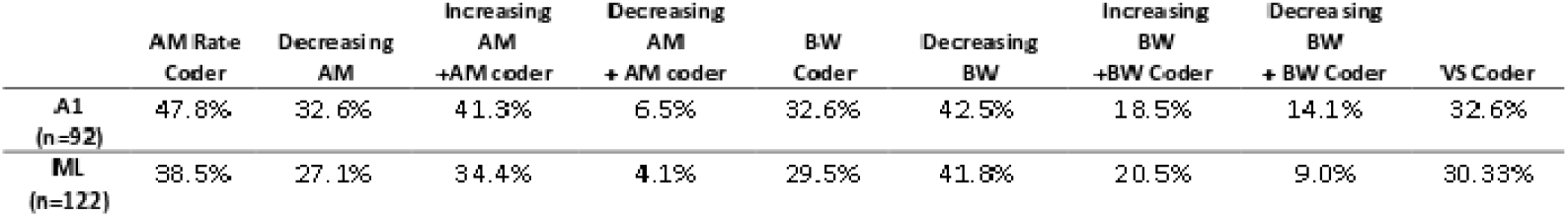

|  | AM Rate<br>Coder | Decreasing<br>AM | Increasing<br>AM<br>+AM coder | Decreasing<br>AM<br>+ AM coder | BW<br>Coder | Decreasing<br>BW | Increasing<br>BW<br>+BW Coder | Decreasing<br>BW<br>+ BW Coder | VS Coder |
| --- | --- | --- | --- | --- | --- | --- | --- | --- | --- |
| **A1**<br>(n=92) | 47.8% | 32.6% | 41.3% | 6.5% | 32.6% | 42.5% | 18.5% | 14.1% | 32.6% |
| **ML**<br>(n=122) | 38.5% | 27.1% | 34.4% | **4.1%** | 29.5% | 41.8% | 20.5% | 9.0% | 30.33% |

A large population of neurons decreased firing rate for increasing AM depth (‘decreasing cells’) in both A1 and ML (Table 1). We also found that nearly half of the neurons in both A1 and ML decreased firing rate for increasing ΔBW (Table 1). However, the population of neurons *that significantly* encoded AM was largely dominated by cells that increased firing rate for increasing AM depth in both A1 and ML, with only 13.6% of AM encoders classified as ‘decreasing’ units in A1, and 10.6% of AM encoders ‘decreasing’ in ML. Among significant ΔBW encoders, the population was more evenly split between ‘increasing’ and ‘decreasing’ units in both A1 and ML: 43.3% of ΔBW encoders have ‘decreasing’ functions in A1 vs. 30.6% in ML. In both A1 and ML, there was a significant positive correlation between AM and BW selectivity, so cells that tended to increase firing rate for increasing AM levels, also tended to increase firing rate for increasing ΔBW levels (For FSI_AM_vs. FSI_BW_, A1 Pearson’s rho = 0.3143, *p* = 0.002; ML Pearson’s rho = 0.3109, *p* = 5.3 e-4) (Figure 1). In this feature selective attention task, we found no significant difference between A1 and ML in the proportions of ‘increasing’ and ‘decreasing’ encoding cells for either AM (‘Increasing’ *p* = 0.21 χ^2^ test; ‘Decreasing’ *p* = 0.11 χ^2^ test) or BW (‘Increasing’ *p* = 0.22 χ^2^ test; ‘Decreasing’ *p* = 0.52 χ^2^ test).

### Vector strength encoding

We found a similar proportion of cells in A1 and ML that significantly phase-locked to AM (p = 0.77, χ^2^ test), as measured by phase-projected vector strength (VS_pp_) (Table 1). As in previous reports (Niwa et al. 2013), we found VS_pp_ to be weaker in ML than A1 (Figure 2, *p* < 0.05 at all AM depths, Wilcoxon rank-sum Test). In both A1 and ML, there was no significant difference in phase-locking (VS_pp_) between the attend AM and attend ΔBW conditions (*p* > 0.05, signed-rank test, Figure 2).

**Figure 2:**
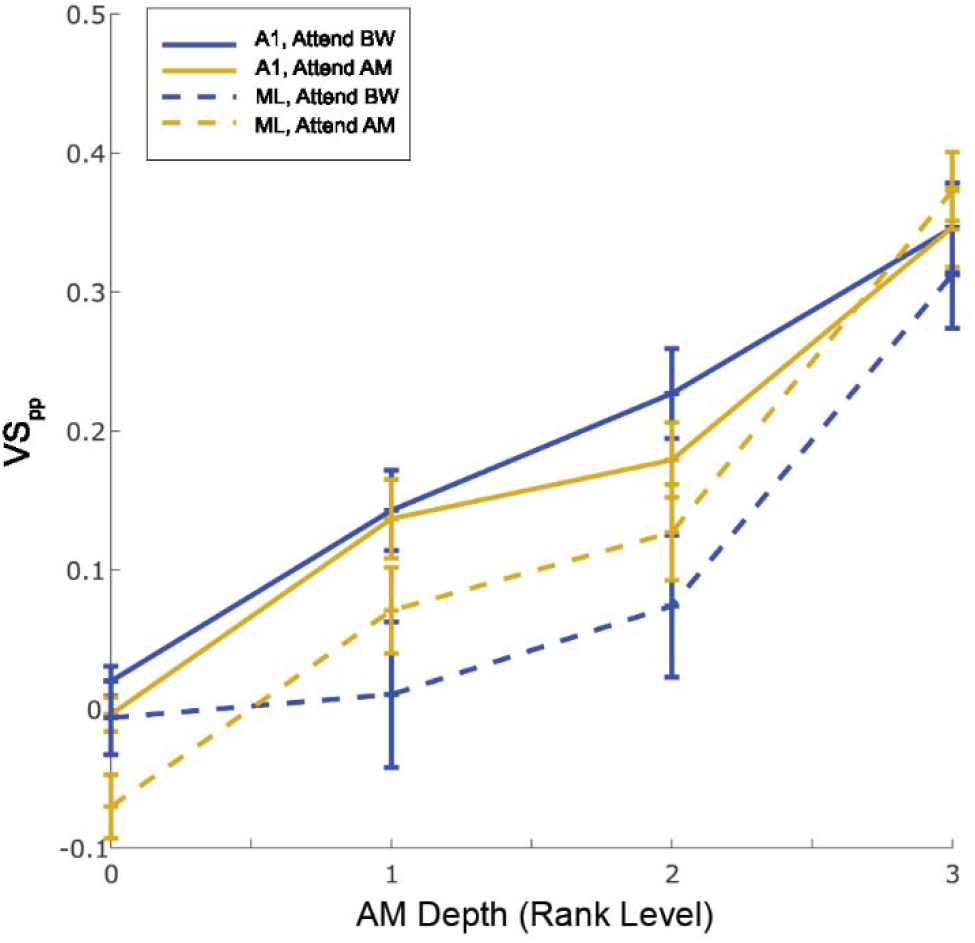
Average phase locking ability of single units in A1 (solid lines) and ML (dashed lines), as measured by phase projected vector strength (VS_pp_). VS_pp_ is greater in A1 (solid) than ML (dashed) at low AM depths (AM level 1, *p* = 0.01; AM level 2, *p* = 0.002, Wilcoxon Ranked Sum), though phase locking is more similar at the highest AM depth (p = 0.73, Wilcoxon ranked sum). There was no significant difference in either area between attend AM (gold) attend ΔBW (blue) conditions, (p > 0.05 for all AM levels, Wilcoxon signed rank test).

### Feature discriminability and context effects

We used the signal detection theory-based area under the receiver operating characteristic (ROCa) to measure how well an ideal observer could detect the presence of each sound feature based on the neural responses (either firing rate or VS_pp_). Increases in the levels of both features tended to yield increasing ROCa (A1 AM Spearman rho = 0.15, BW Spearman rho =.06; ML AM Spearman rho = 0.13, BW Spearman rho = 0.05) (Figure 3). However, there was no significant effect of attentional condition on either feature at any level of feature modulation for either A1 (Figure 3a,c) or ML (Figure 3b,d).

**Figure 3:**
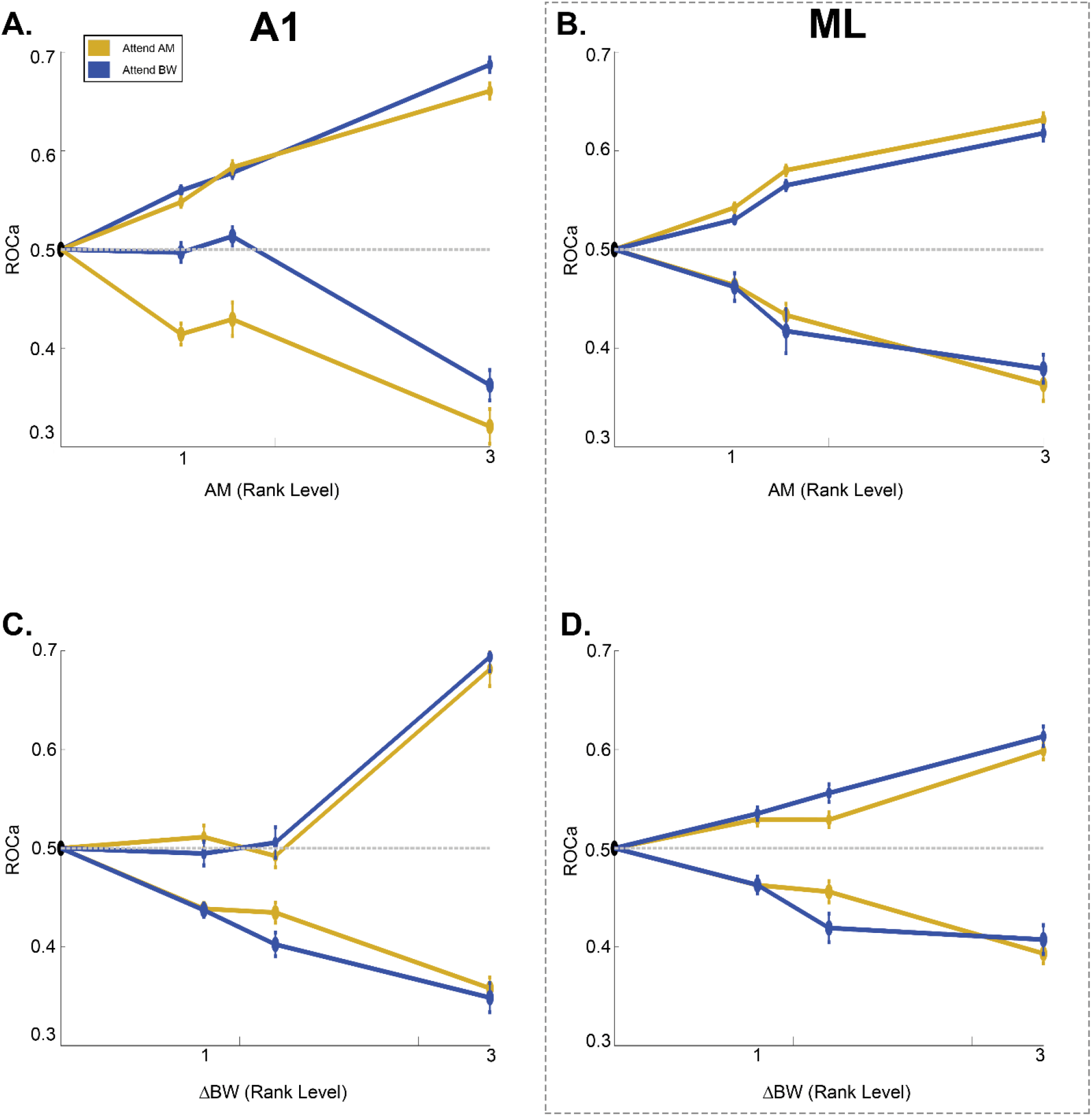
Firing rate based ROCa for each feature by attention condition. Blue lines indicate attend BW condition, yellow lines indicate attend AM condition. A. AM encoding in A1 (38 increasing, 6 decreasing cells). B. AM encoding in ML (42 increasing, 5 decreasing cells) C. BW encoding in A1 (17 increasing, 13 decreasing cells) D. BW encoding in ML (25 increasing, 11 decreasing cells). There was no significant effect of attentional condition on either feature at any level of feature modulation for either A1 **(A,C)** or ML **(B,D).**

VS_pp_ -based discrimination (ROCa) of AM from unmodulated sounds was better at the lowest modulation depth in A1 than in ML (p = 0.02, Wilcoxon Rank Sum Test, Figure 4). At the higher modulation depths, VS_pp_ -based discrimination was similar in A1 and ML (p = 0.99 AM depth 2; p = 0.26, AM depth 3; Wilcoxon Rank Sum Test; Figure 4). However, there was no significant difference in VS_pp_ discriminability between attention conditions for any modulation depth (p > 0.05, signed-rank test, Figure 4).

**Figure 4:**
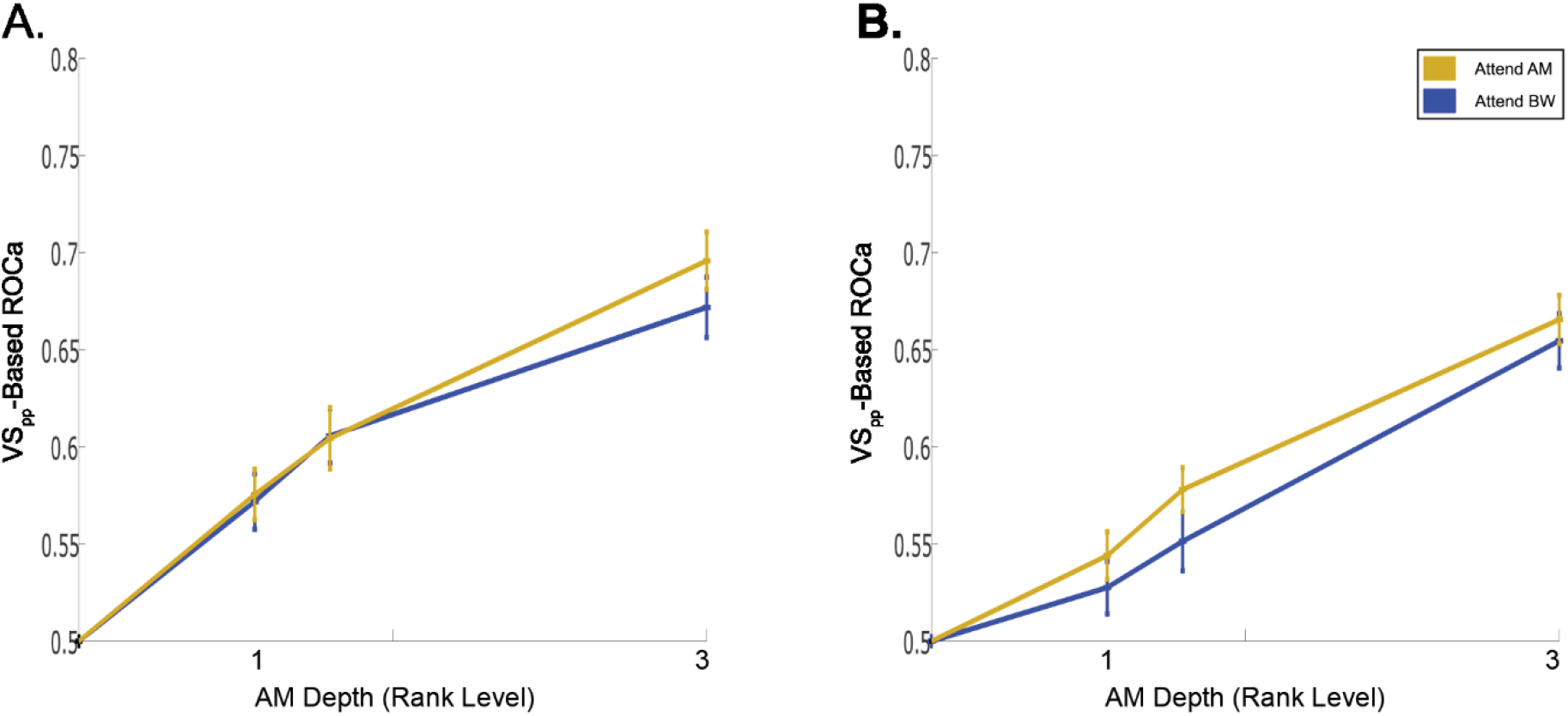
VS_pp_-based discriminability (ROCa) of AM from unmodulated sounds in A1 and ML for attend AM (yellow) and attend ΔBW (blue) conditions. **A.** In Al, VS_pp_-based discriminability of AM is not significantly different between attention conditions (p > 0.05, signed-rank test) B. In ML, VS_pp_-based ROCa does not differ between attentional conditions (p > 0.05, signed-rank test). At low modulation depths (AM depth rank = 1). A1 had significantly better AM discriminability than ML (p = 0.02, rank sum test), however they were not significantly different at the higher modulation depths (AM ranks 2 and 3, p > 0.05, rank sum test).

### Choice-related activity

We found a similar proportion of neurons in A1 (19.5%) and ML (26.2%) with significant choice-related activity during the stimulus window (p = 0.31, χ2 test). In both areas, the population of neurons with significant choice-related activity during the response window (from S2 end to joystick movement) was larger than during the stimulus window, and the proportions of neurons were again similar between the two areas (41.3% A1, 34.4% ML, *p* = 0.41, χ2 test).

In A1, during the attend AM condition, CP values were evenly distributed about 0.5 during both the stimulus presentation (S2 median CP = 0.50, *p* = 0.87 signed-rank test) and the response window (RW median CP = 0.49, *p* = 0.43 signed-rank test) (Figure 5a,c). In contrast, during the attend ΔBW context, the CP values tended to be lower than 0.5 during both the stimulus (S2 median CP = 0.49, *p* = 0.02 signed-rank test) and the response window (RW median CP =0.46, *p* = 4.2 e-8 signed-rank test) (Figure 5b,d). That is, during the attend ΔBW condition, the population of neurons tended to decrease firing rate when reporting target feature detection, whereas during the attend AM condition, it was equally likely for a neuron to increase firing rate for a report of target detection as it was for a report of target absence. There was a significant difference in the population CP distributions between attention conditions during the RW (Attend AM median = 0.49, Attend ΔBW median = 0.46, *p* = 0.004, signed-rank test), though not during the S2 (p = 0.06, signed-rank test).

**Figure 5:**
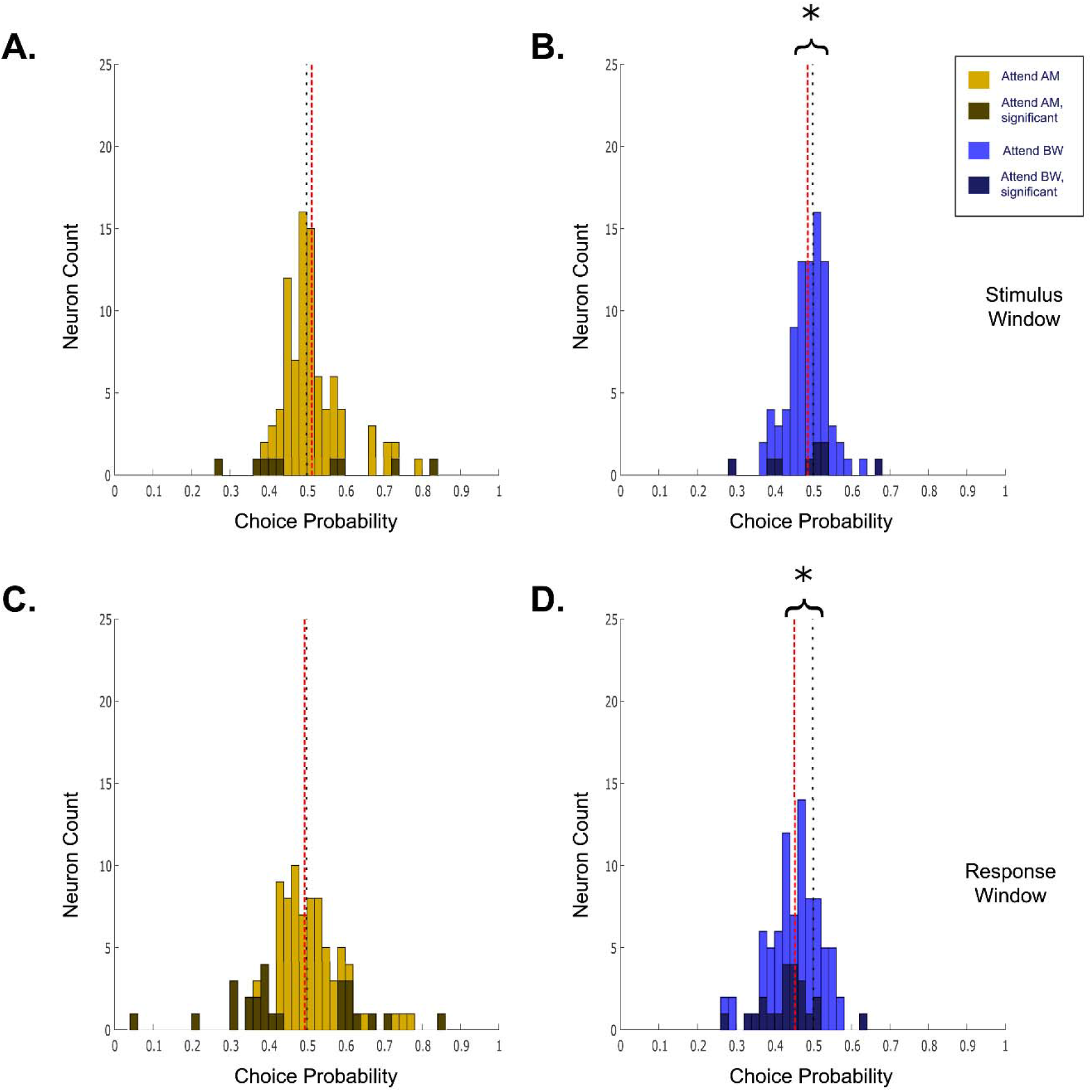
Choice probability in Al. Values closer to 0 indicate increased activity for ‘feature-absent’ response, whereas 1 indicates increased activity for ‘feature-present’ response. Darker colored bars indicate cells with significant choice activity. Black dotted line indicates 0.5, red dashed line denotes the population median. CP during the attend AM condition is evenly distributed about 0.5 in both A. the stimulus window (median = 0.50, *p* = 0.87, signed-rank test) and C. the response window (median = 0.49, *p* = 0.43, signed-rank test). In the attend ΔBW condition, CP values tended to be less than 0.5 in both B. the stimulus window (median = 0.49, *p* = 0.02 signed-rank test) and D. the response window (median = 0.46, *p* = 4.2 e-8 signed-rank test). There was a significant difference in the population CP distributions between attention conditions during the RW (p = 0.004, signed-rank test), though not during the S2 (p = 0.06, signed-rank test).

The choice-related activity in ML was similar to that reported above in A1 during the S2. During the attend AM condition, activity was evenly distributed about 0.5 (S2 median CP = 0.50, p = 0.94, signed-rank test) (Figure 6a). During the attend ΔBW condition, average CP was less than 0.5 (S2 median CP = 0.49, p = 0.043 signed-rank test) (Figure 6b). However, during the response window, CP values were less than 0.5 in both the attend AM condition (median CP = 0.48, p = 0.004 signed-rank test) and the attend ΔBW condition (median CP = 0.47, p = 2.7 e-5 signed-rank test) (Figure 6c,d). This is in contrast to A1 where CP values tended to be lower than 0.5 only in the attend BW condition. There was no significant difference in the distribution of CP values in ML neurons between the attend AM and attend ΔBW conditions during the S2 (p = 0.15, signed-rank test). However, there was a significant difference in the CP distribution between the attend AM and attend ΔBW conditions in ML during the response window (Attend AM median = 0.48, Attend ΔBW median = 0.47, p = 0.033 signed-rank test), reflecting the population shift to CP values less than 0.5 in the attend ΔBW condition compared to the attend AM condition.

**Figure 6:**
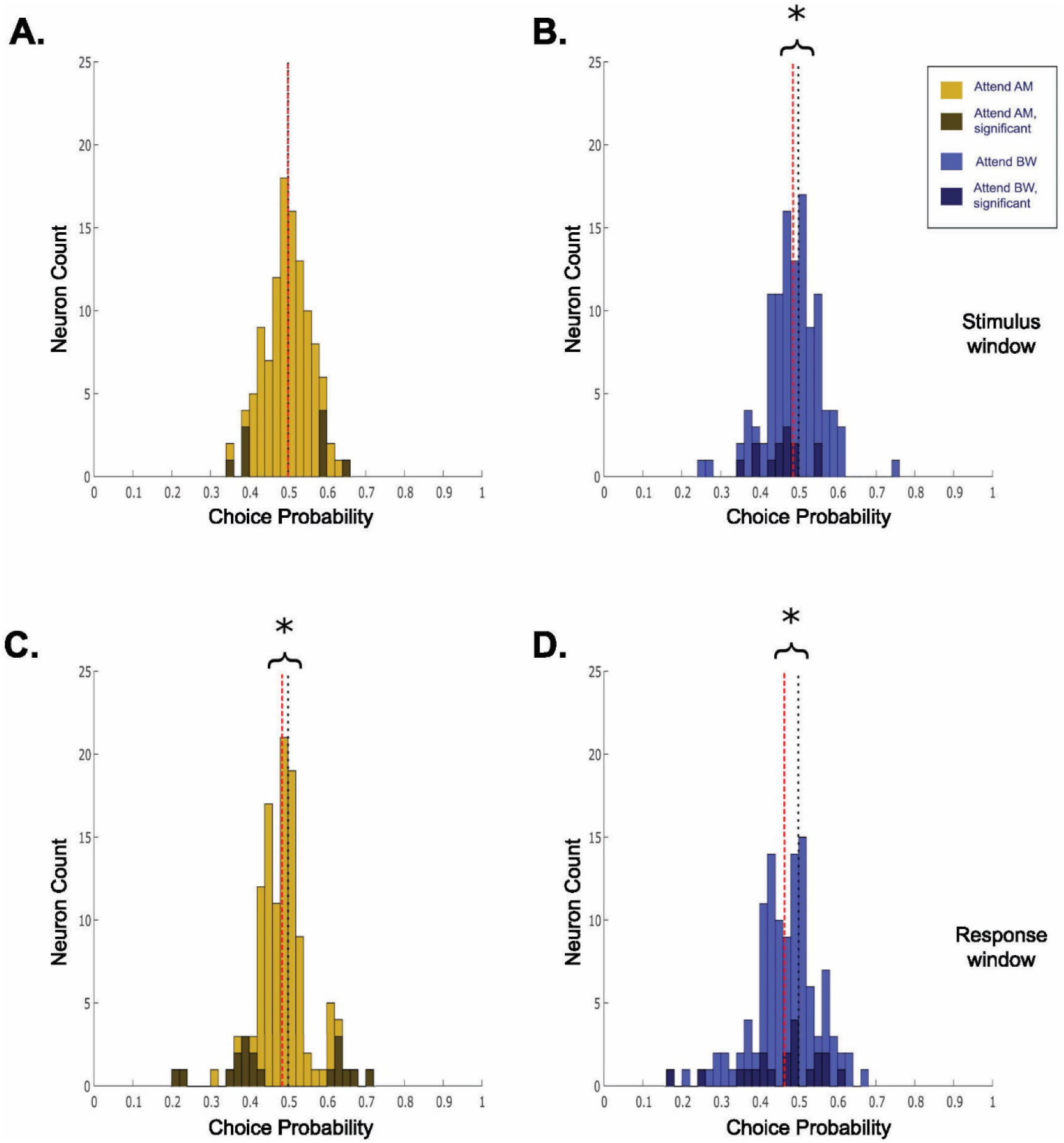
Choice probability in ML, as in Figure 5. CP during the attend AM condition is evenly distributed about 0.5 in A. the stimulus window (median = 0.50, *p* = 0.94, signed-rank test). However, in the response window C. CP values tended to be less than 0.5 (median = 0.48, *p* = 0.004, signed-rank test). In the attend ΔBW condition, CP values tended to be less than 0.5 in both B. the stimulus window (median = 0.49, *p* = 0.043 signed-rank test) and D. the response window (median = 0.47, *p* = 2.7 e-5 signed-rank test). As in Al, there was a significant difference in the population CP distributions between attention conditions during the RW (p = 0.033, signed-rank test), though not during the S2 (p = 0.15, signed-rank test).

## Discussion

We found a large proportion of cells in ML that decreased firing rate with increasing AM detectability, similar to previous findings in ML (Johnson et al. 2020; Niwa et al. 2013). However, unlike these previous studies where ML had a significantly larger population of cells with decreasing AM depth functions than A1, we found a similar proportion of A1 neurons with decreasing AM depth functions. Further, the majority of neurons in both A1 and ML significantly encoding AM depth had increasing AM depth functions. This suggests that the encoding of amplitude modulation can be flexible depending upon the behavioral and sensory demands of the task. In essence, with increased perceptual difficulty, stimulus/feature ambiguity, and task difficulty it may be necessary for A1 to develop a more robust and appropriate code in order to solve the task, and for ML to take on more of the sensory processing, and thus the encoding schemes look more similar between these two areas.

We also found a large population of cells in both A1 and ML that were sensitive to changes in bandwidth. This was particularly surprising as the changes in bandwidth were relatively small compared to the 9-octave wide unmodulated noise. It’s possible that the ΔBW encoding we saw was due to an increasing concentration of power in the middle frequencies of the broadband noise as the level of bandwidth restriction increased. It could also be caused by decreasing power in flanking inhibitory bands. Further studies investigating if and how neurons in A1 and ML encode small changes in spectral bandwidth to broad-band sounds under power-matched conditions could be enlightening.

Using phase-projected vector strength (VS_pp_) as a measure of temporal coding, neither ML nor A1 single neurons showed attention-related changes in VS_pp_-based sensitivity to AM or VS-based choice-related activity. This is consistent with previous results from our lab showing smaller effects for VS_pp_-based attention and choice than for firing rate (Niwa et al. 2013). A recent study that could help interpretation of this result shows thalamic projections to the striatum (an area involved in decisions and possibly attention) relay information about temporally modulated sounds in the form of phaselocking, whereas cortical projections to the striatum only convey information about temporally modulated sounds with average firing rate over the stimulus (Ponvert and Jaramillo 2019).

Attending to the target-feature did not significantly improve single neuron amplitude modulation or bandwidth restriction detection in A1 or ML. This seems surprising considering the wide array of effects that have been previously reported in auditory cortex related to different tasks, and behavioral contexts (Atiani et al. 2014; Bagur et al. 2018; Buran et al. 2014; Francis et al. 2018a; Niwa et al. 2012b; Otazu et al. 2009; Lakatos et al. 2013; Angeloni and Geffen 2018; Sutter and Shamma 2011). In macaque monkeys, an improvement in both rate-based and temporal AM encoding was observed in A1 and ML neurons when animals performed a single-feature AM detection task compared to when animals passively listened to the same stimuli (Niwa et al. 2013, 2015). We did not see a similar level of encoding improvement, possibly due to the more fine-tuned form of attention needed to perform this task.

One might expect to observe smaller effects from this more selective form of attention than in a passive versus active listening task, as the difference between attending to one feature of a sound compared to another is much smaller than switching between paying attention to a sound and passive sound presentation. Furthermore, arousal, as measured with pupillometry, has recently been shown to correlate with increases in activity, gain and trial-to-trial reliability of A1 neurons (Schwartz et al. 2019), which could account for some of the effects seen in task engagement paradigms.

Feature-based attention has been shown to have gain effects on neurons tuned to the attended feature in visual cortex (Ni and Maunsell 2019; Treue and Trujillo 1999). It is possible that we did not see a similar gain effect of feature attention in AC due to the mixed-selectivity we and others (Chambers et al. 2014) found in the encoding of these features (i.e. most neurons are sensitive to both AM and ΔBW). However, it is likely that mixed-selectivity is not the only reason we did not see a gain effect. In a study where rats performed a frequency categorization task with shifting boundaries, Jaramillo and colleagues similarly found that neurons in AC did not improve their discriminability with attentional context (Jaramillo et al. 2014). This similar lack of enhancement seen in a task where only a single feature is modulated, suggests that the mechanism for feature attention in auditory cortex could be enacted via a different mechanism.

In visual cortical studies probing *selective* feature attention – where the subject must distinguish between features within a single object, rather than object-or place-oriented, feature-based attention – results have been similarly complex. At the level of the single neuron, there have not been clear, gain-like improvements in the sensitivity to the attended feature (Chen et al. 2012; Mirabella et al. 2007; Sasaki and Uka 2009; Uka et al. 2012). Further, the effects of feature-selective attention seem to be dependent upon not just the tuning preferences of a neuron, but also the strength of its tuning (Ruff and Born 2015). These studies, along with our own, suggest that segregation of features within an object may require a different mechanism relative to object-directed, feature-based attention.

In each of the feature-selective attention studies cited above, a common observation is that single neurons in sensory cortex have mixed selectivity for the features in the task, as opposed to being uniquely responsive to one feature or another. Such mixed selectivity among single neurons may permit sophisticated, flexible computations at the population level (Fusi et al. 2016). It thus seems likely the mechanism for feature-selective attention lies not at the level of the single neuron, but rather requires the integration of activity from a larger population of neurons. A feature-selective study using ERPs found that the neural responses to identical stimuli varied when the subjects attend to different features of the stimulus (Nobre et al. 2006). The single neuron and neural circuit mechanisms underlying this effect remain unclear. One such possible mechanism might be the structure of correlated variability within the population, which has been shown to be modulated by feature-selective attention (Downer et al. 2017b). Another study, simulating populations by pooling single-neurons across A1 recordings permitted clear segregation of these two features, as well as an enhancement in discrimination of the attended feature (Downer et al. 2020). Further studies investigating feature-selective attention at the level of populations of neurons are necessary to better understand the underlying mechanisms.

We did see an interesting difference in the distribution of choice-related activity between the attentional conditions, where the correlation between firing rate and choice shifted direction between conditions. During the attend AM context, CP was evenly distributed about 0.5 with some neurons showing significant choice activity at either extreme. In contrast, during the attend BW context, CP values were shifted towards 0, with very few neurons having significant choice-related activity greater than 0.5 (increasing firing rate for ‘feature-present’ response). Neurons in auditory cortical areas may also modulate their responses to motor events (Brosch 2005). Some previous reports on choice-related activity have been difficult to interpret, as they employed a Go/No-Go task format in which one perceptual choice required a movement and the other choice did not (Brosch 2005; Niwa et al. 2013). Therefore, the choice-related activity observed was difficult to disentangle from a general preparation to move. The task reported here was a Yes/No forced-choice task, requiring a motor response to each decision (target present versus target absent). The shift in choice-related activity between attention conditions observed in this force choice task, and another recent study (Guo et al. 2019) shows that this choice-related activity cannot simply reflect motor preparation or action. This then strengthens the possible relationship between this activity and the decision or attention process.

The lack of clear attentional improvement of single neuron feature encoding found in this study suggests one or more of the following: (1) the feature-selective attention required in this task is not implemented at the level of an individual neuron in A1 or ML; (2) the feature-selective attention necessary for this particular task occurs at a later stage in auditory processing; (3) the mixed selectivity of single neurons in A1 and ML for these features complicates the interpretability of the effects of attention at the single neuron level, in contrast to feature-based attention neurons studied found in visual cortex (Martinez-Trujillo and Treue 2004; Maunsell 2015; Maunsell and Treue 2006). While we did not see robust differences in encoding between attentional conditions, the difference in attentional choice-related activity reveals that it is not simply reflective of motor preparation, and suggests that activity correlated to reported choice as early as A1 could be informing perceptual and decision processes.

## Acknowledgements

This work was funded by NIH NIDCD grant DC002514 (MLS), NSF GRFP 1148897 (JDD) and ARCS Foundation Fellowship (JDD)

